## Supplementary figures and images for "A new phylodynamic model of *Mycobacterium bovis* transmission in a multi-host system uncovers the role of the unobserved reservoir"

### Supplementary figure 1

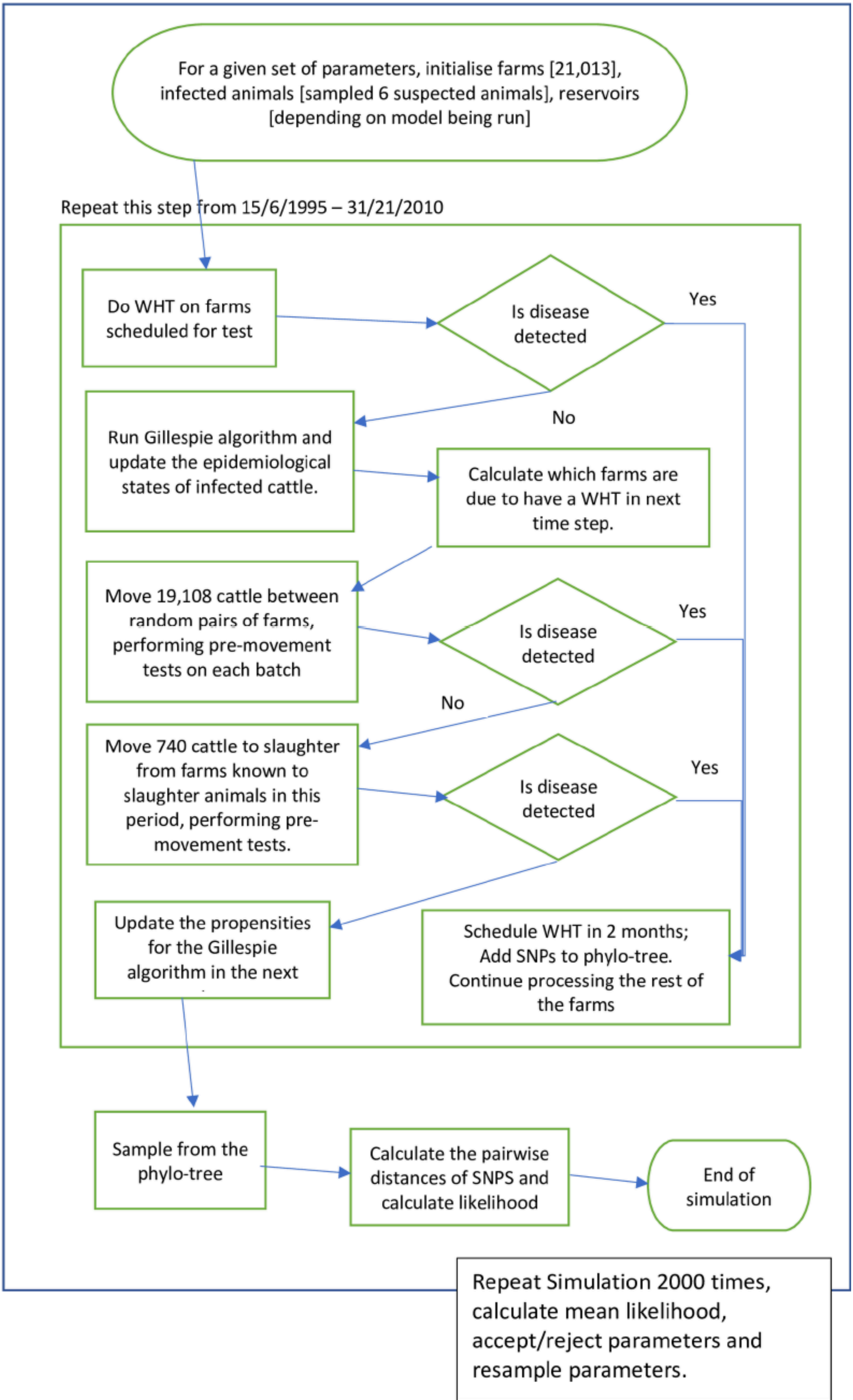

### Supplementary figure 2

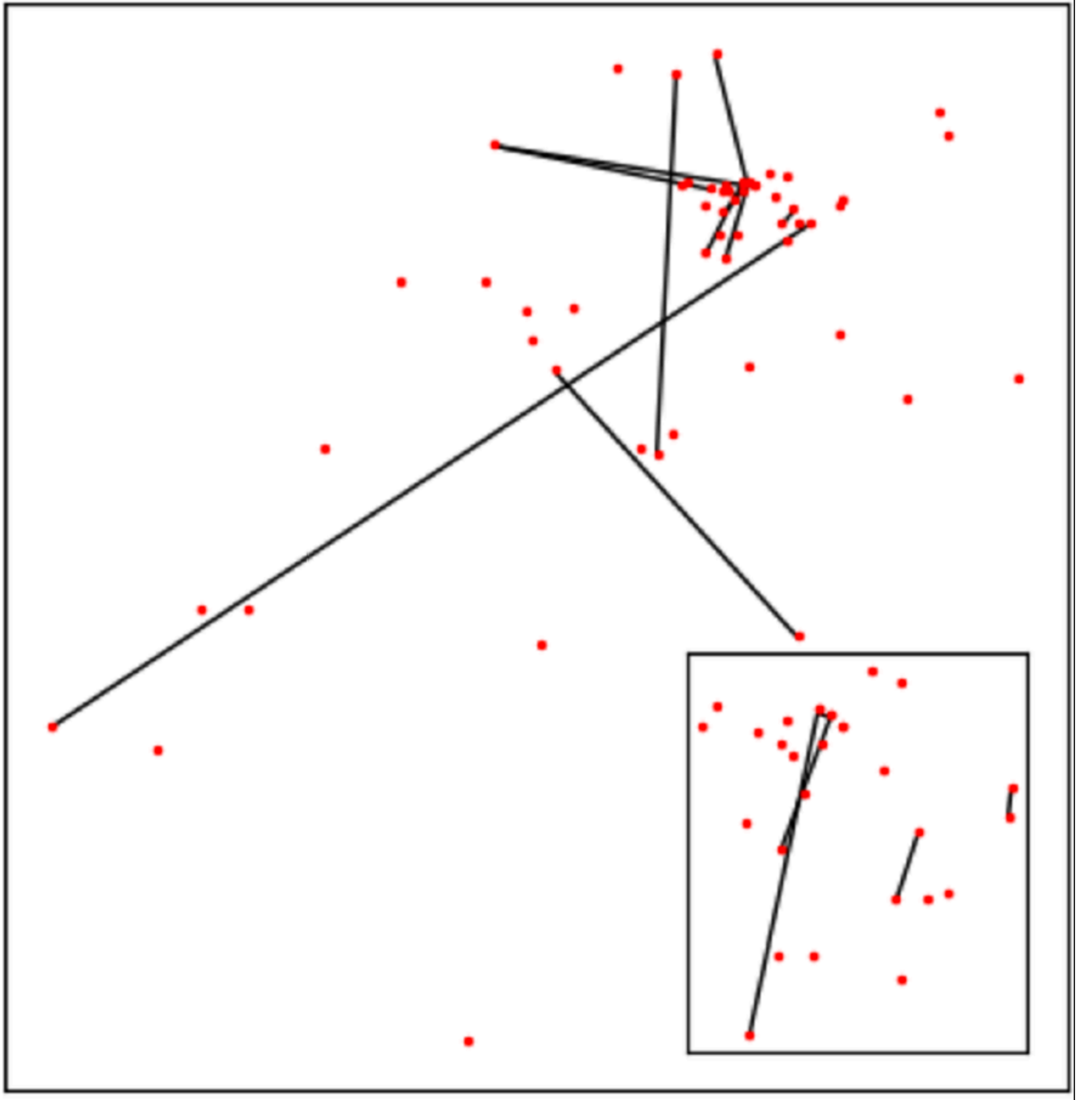

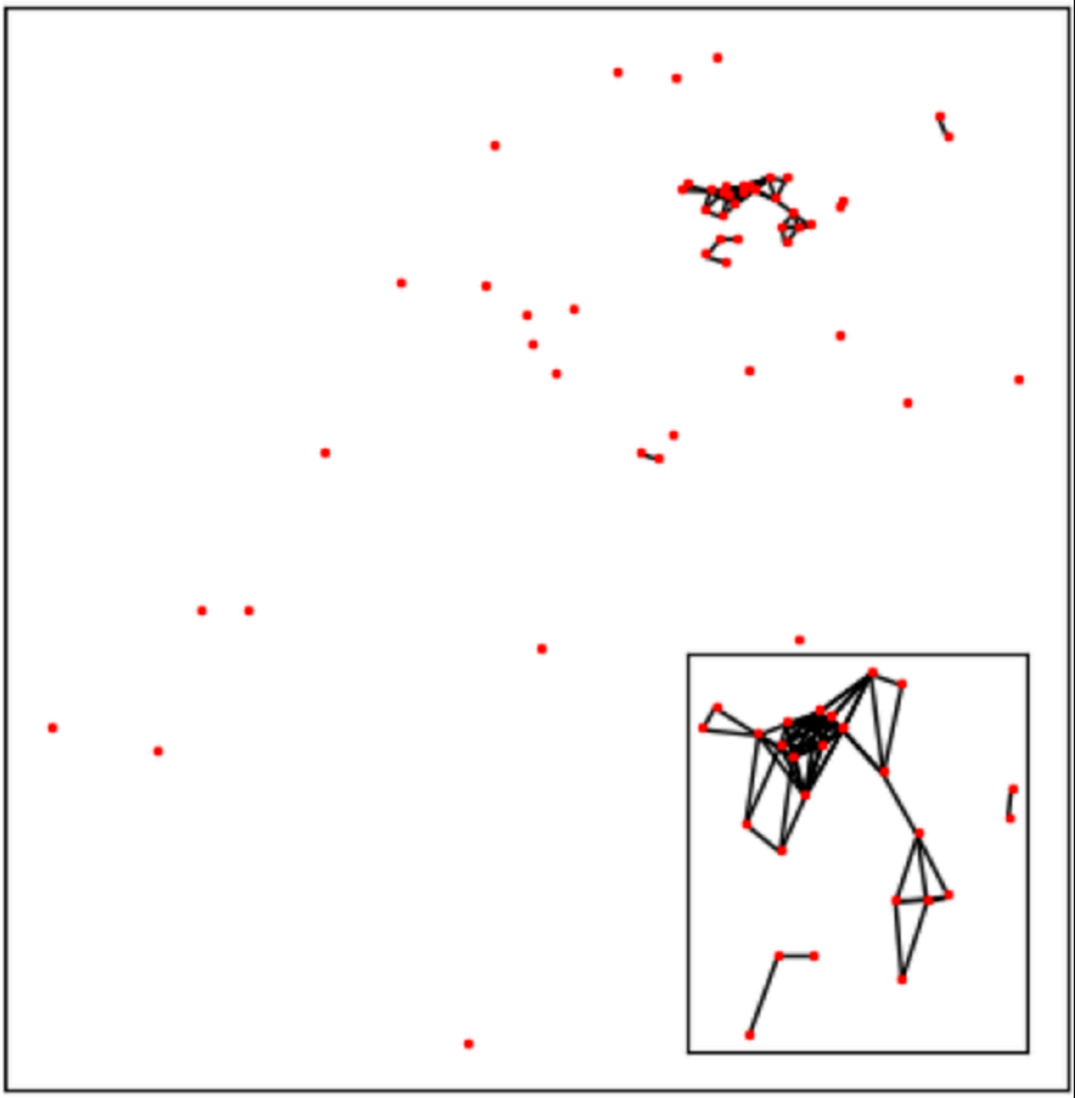

### Supplementary figure 3

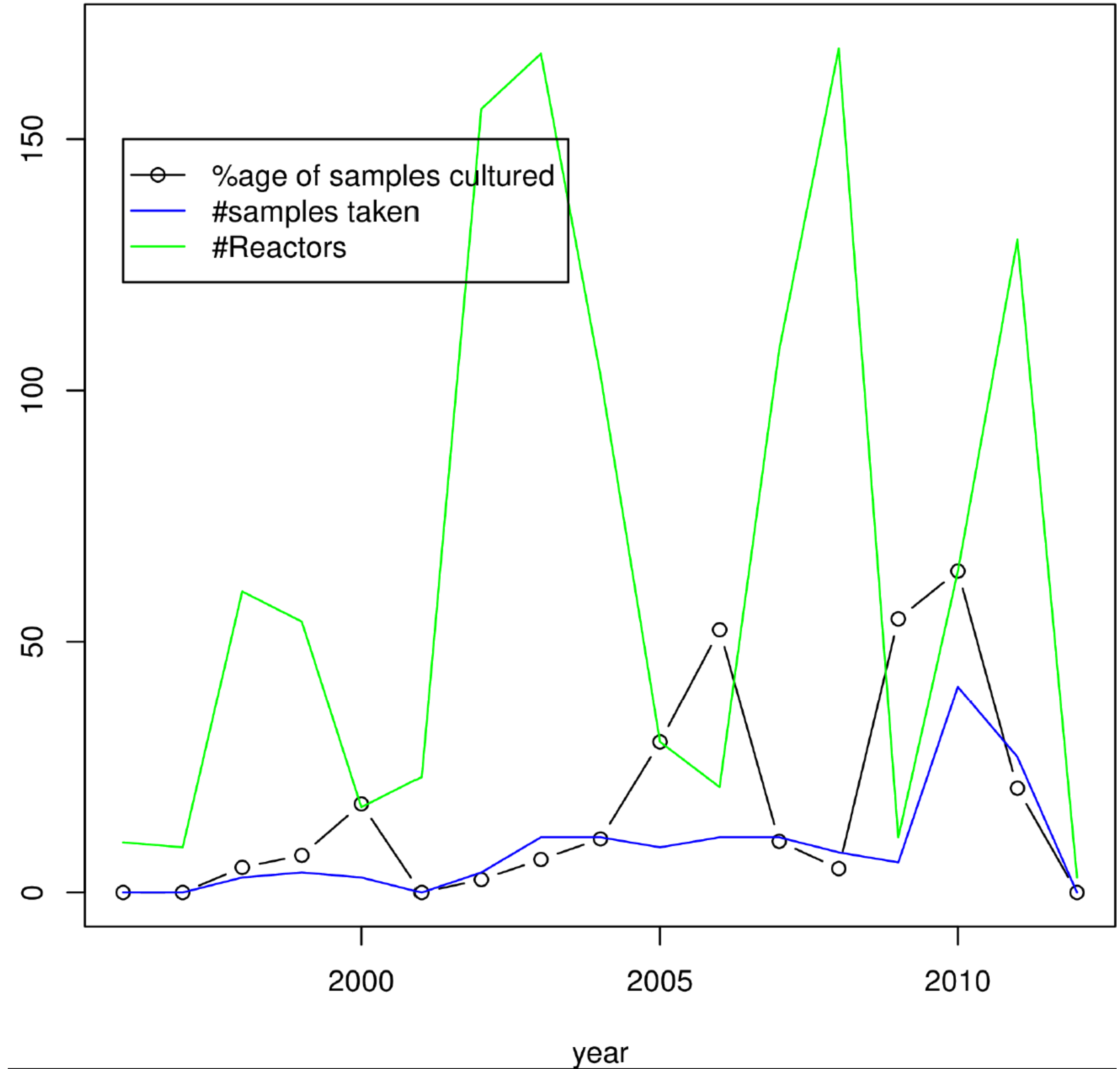

### Supplementary figure 4

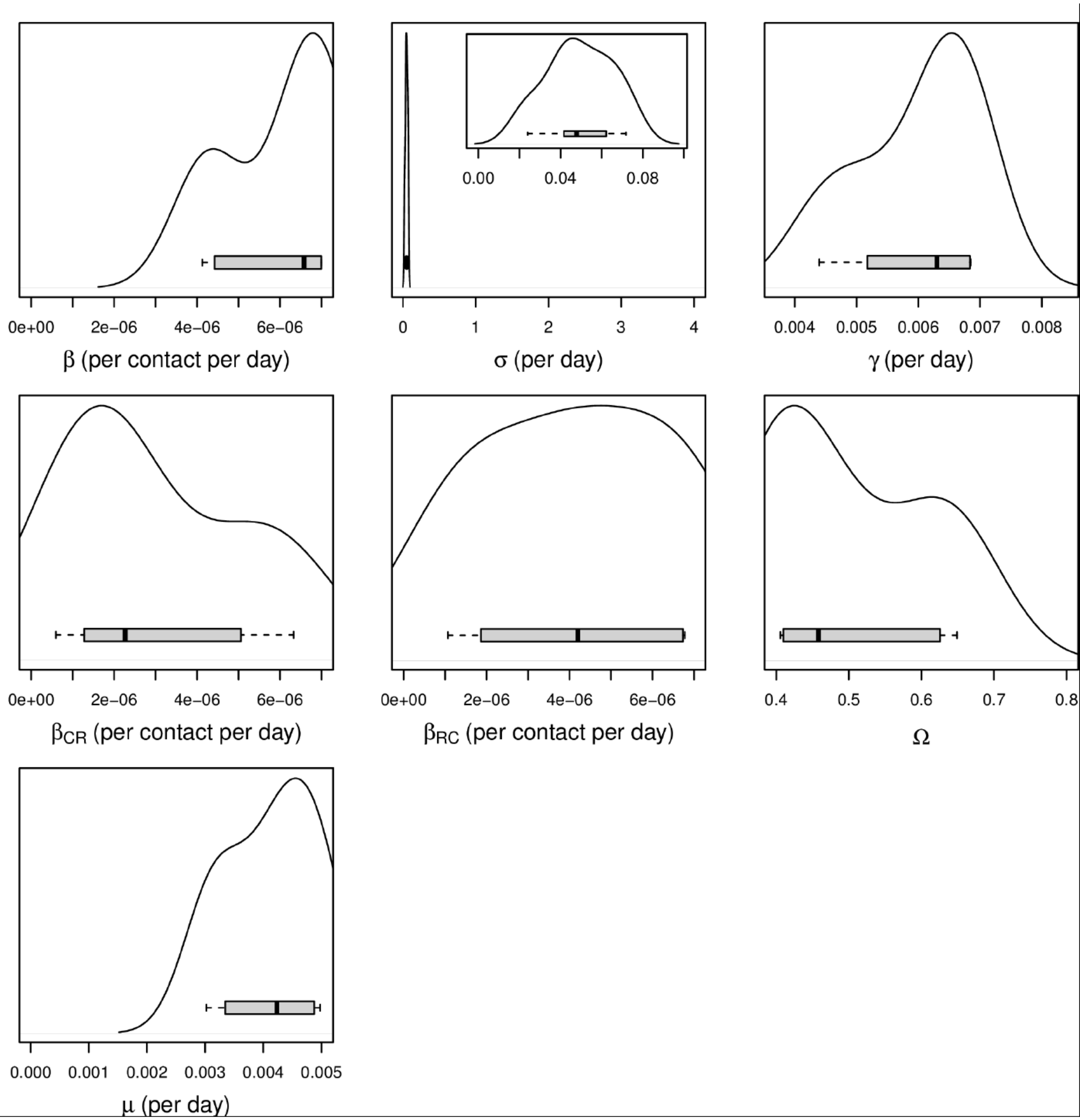

### Supplementary figure 5

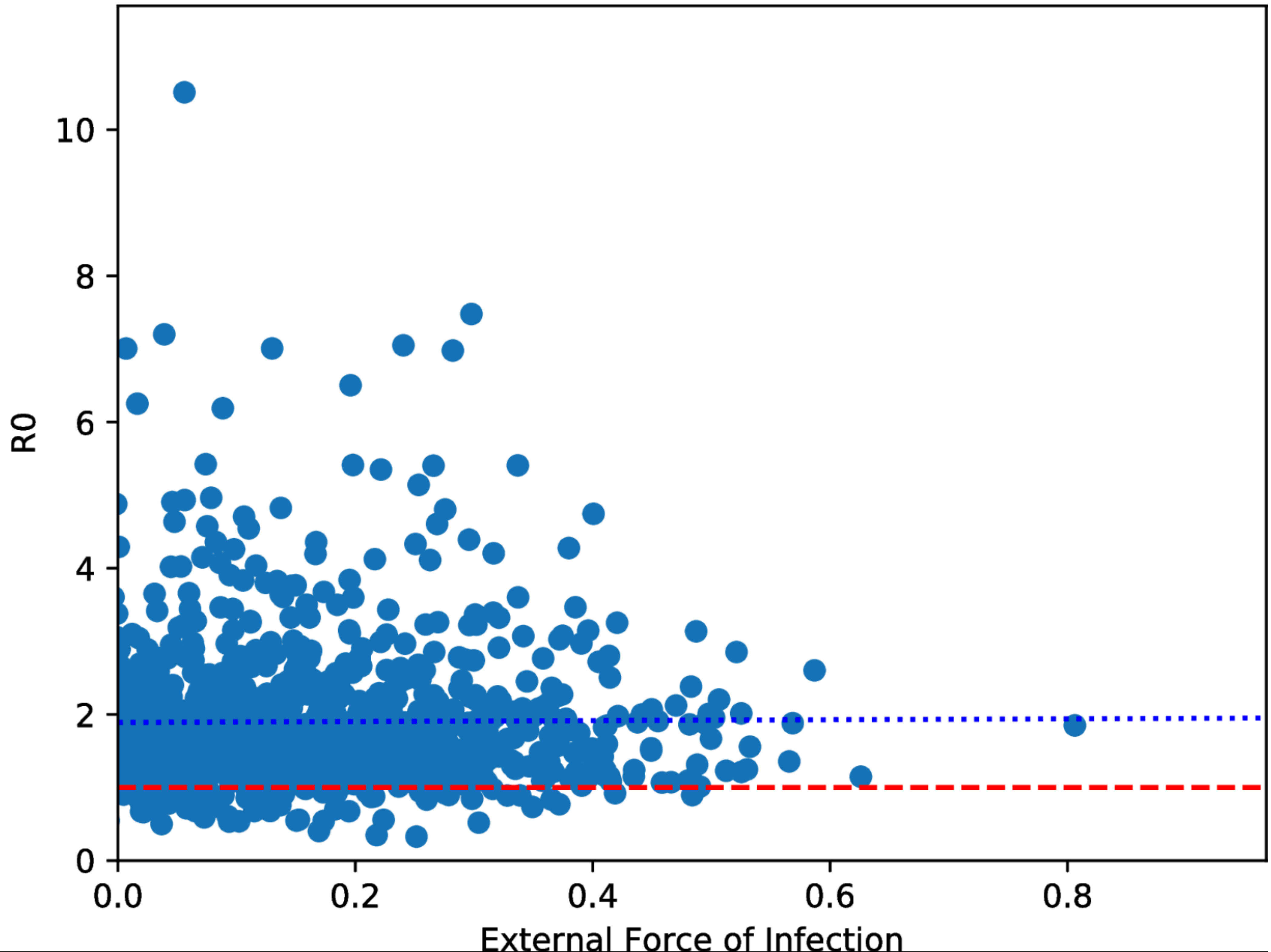
